# Directed evolution of CRISPR-Cas9 to increase its specificity

**DOI:** 10.1101/237040

**Authors:** Jungjoon K. Lee, Euihwan Jeong, Joonsun Lee, Minhee Jung, Eunji Shin, Young-hoon Kim, Kangin Lee, Daesik Kim, Seokjoong Kim, Jin-Soo Kim

## Abstract

The use of CRISPR-Cas9 as a therapeutic reagent is hampered by its off-target effects. Although rationally designed S. pyogenes Cas9 (SpCas9) variants that display higher specificities than the wild-type SpCas9 protein are available, these attenuated Cas9 variants are often poorly efficient in human cells. Here, we have used a directed evolution approach in E. coli to obtain Sniper-Cas9, which shows high specificities without sacrificing on-target activities in human cells.

## Main text

The determination of the Cas9 crystal structure^1^ enabled scientists to rationally design mutant Cas9 proteins (enhanced SpCas9 (eSpCas9) and Cas9-High Fidelity (Cas9-HF)) with higher specificities than wild-type Cas9 (WT-Cas9)^2, 3^. Their design was based on the hypothesis that weakening non-specific interactions between a Cas9-RNA complex and its substrate DNA would reduce both on-target and off-target activities alike. Since on-target activity is generally much higher than off-target activity, these mutant Cas9 variants would show higher specificities than WT while retaining on-target activities. However, many groups have since reported that both eSpCas9 and Cas9-HF were poorly active at many targets tested^4–6^, calling for alternative approaches to improve Cas9 specificity.

We reasoned that directed evolution of Cas9 in E. coli could lead to Cas9 variants with high specificity without sacrificing on-target activities. The system we used consists of E. coli strain BW25141 and a plasmid containing the lethal *ccdB* gene^7–9^ and the Cas9 target sequence: The disruption of the *ccdB* gene by Cas9-mediated plasmid DNA cleavage is essential for cell survival, creating a positive selection pressure. In addition, a Cas9 off-target sequence that differs from the on-target sequence by a few mismatches is introduced in the E. coli genomic DNA: Double strand breaks (DSBs) in E. coli genomic DNA lead to cell death. We combined such negative selection pressure with *ccdB* plasmid-based positive selection pressure to develop ‘Sniper-screen’, which selects for Cas9 variants with increased specificities. Note that Cas9 variants with poor on-target activities or poor specificities cannot survive in this selection system.

We first inserted a 500-bp PCR product containing an *EMX1* fragment into the genomic DNA of the BW25141 strain using a protocol involving transposase^10^ (Figure 1a and Supplementary figure 1). The resulting BW25141-EMX1 strain was transformed with two plasmids (Supplementary figure 2): a plasmid expressing *ccdB* under the control of the pBAD promoter, which is induced by arabinose, and a plasmid expressing an sgRNA under the control of the pltetO1 promoter, which is induced by anhydrotetracycline (ATC). A target sequence with mismatches relative to the *EMX1* site was inserted into the *ccdB* plasmid and the matching guide sequence was inserted into the sgRNA plasmid. In a Sniper screen, the resulting E. coli strain is electroporated with a pooled library expressing mutant Cas9 variants under the control of a CMV-pltetO1 dual promoter induced by ATC. There are four possible outcomes with respect to DNA cleavage in the *ccdB* plasmid and the genomic DNA (Fig. 1a): Only a mutant variant of Cas9 that discriminates the on-target sequence present in the *ccdB* plasmid from the off-target sequence present in the E. coli genomic DNA can survive. Our system uses separate Cas9 and sgRNA plasmids, making it easy to change sgRNA-encoding and target sequence-containing plasmids in subsequent rounds of selection. Because Cas9 can also be expressed in mammalian cells via the CMV promoter, it is possible to check the on-target and off-target activity of the pool obtained in each round. In addition, Cas9 and sgRNA expression are controlled by the pltetO1 promoter, allowing regulation of gene expression by up to 5000-fold^11^. By increasing the concentration of ATC, the screening conditions become more lenient for *ccdB* cleaving positive selection and harsher for genomic DNA cleaving negative selection. Such adjustments were necessary to find the optimum conditions at which control experiments using WT-Cas9 and a null vector showed a large window for cleavage for each target-sgRNA pair (Supplementary figure 3).

**Figure 1.**
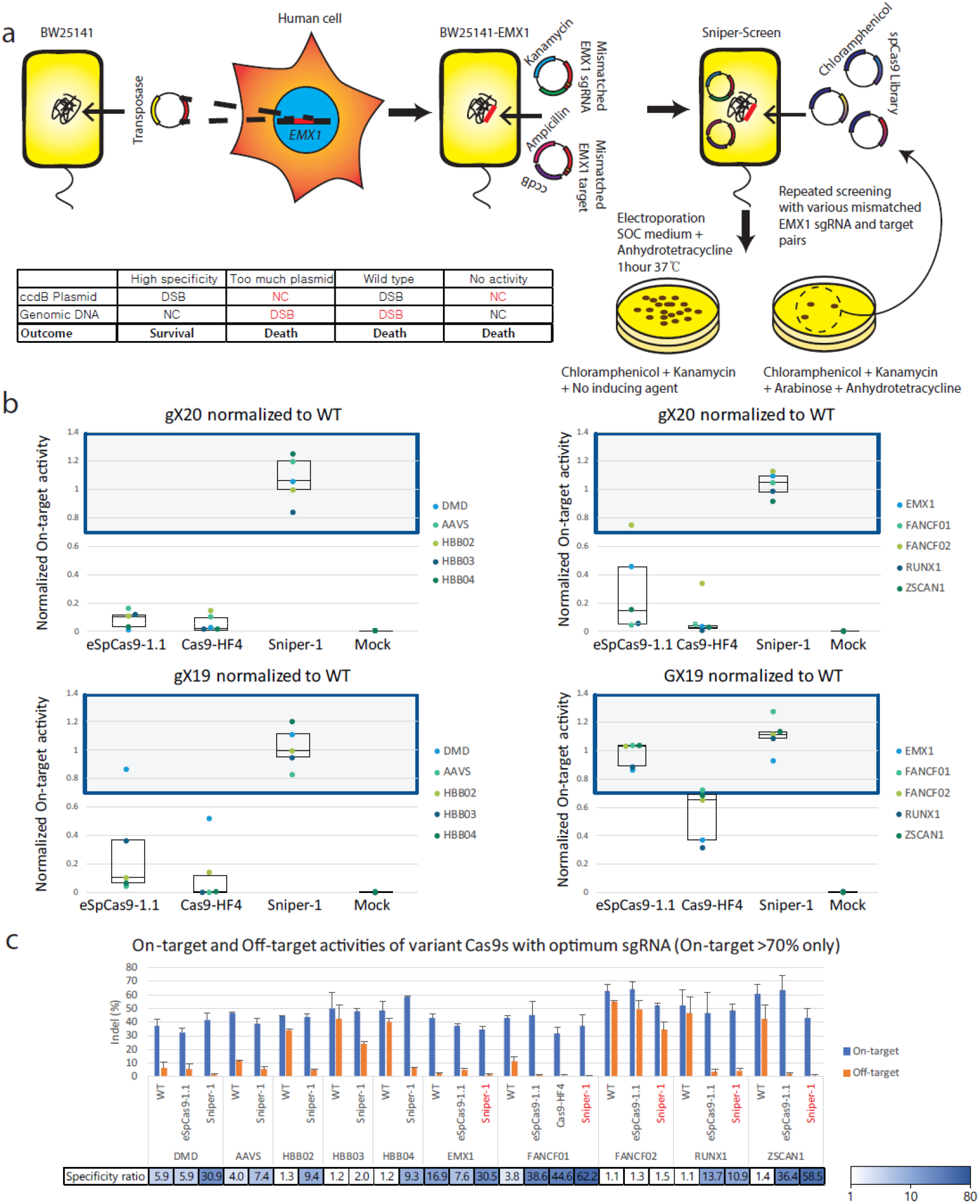
(**a**) Schematic of the Sniper-screen. DSB, double strand break; NC, no cleavage. Cell death is indicated by red type. (**b**) Box plots of on-target activities normalized to WT-Cas9 activity. The boxes represent the 1^st^, 2^nd^ and 3^rd^ quartiles of normalized on-target activities of eSpCas9-1.1, Cas9-HF4, Sniper-1 and mock. The grey area represents on-target activities higher than 70% of the WT-Cas9 activity. The 5’ guanine of the sgRNA may (GX19) or may not (gX19 or gX20) match the target. Error bars were omitted for simplicity (n=3) (c) On-target and off-target activities of Cas9 variant-sgRNA combinations with more than 70% on-target activity relative to WT-Cas9. Specificity ratios were determined by dividing the on-target activity by the off-target activity. Results with gX19 or GX19 sgRNAs are shown in black type and those with gX20 sgRNAs are shown in red type. Error bars indicate s.e.m. (*n*=3)

SpCas9 mutant libraries with random errors in the whole Cas9 sequence were constructed using three different kits, resulting in library complexities of up to 10^7^ overall. Two independent sets of screenings were performed using the libraries (Supplementary figure 4). The first set started with more lenient screening conditions and progressed toward more stringent conditions: DNA shuffling was performed in the middle of the process to enrich the diversity. The second set employed harsh conditions without DNA shuffling. After the final selection steps for both screening sets, the pooled libraries were tested in mammalian cells to measure the specificities of the Cas9 variants compared to WT-Cas9 (Supplementary figure 5). The pooled libraries showed higher specificities than WT-Cas9; the first set was more specific than the other set. One hundred colonies were picked from both sets and the Cas9-encoding DNA sequences were fully sequenced. Three Cas9 variants were identified from the first set, which were designated Sniper-1, Sniper-2 and Sniper-3 (Supplementary figure 15). The mutations were dispersed throughout domains revealed in the crystal structure^1^; none of them had previously been reported in rational design studies (Supplementary figure 6). Site-directed mutagenesis analyses revealed that no single point mutation drastically improved the specificity of WT-Cas9 (Supplementary figure 7). Since we performed DNA shuffling reaction in the middle of our screening, we assumed that the mutant with the best combination of point mutations survived in our screening.

Among the three different variants, Sniper-1 showed the highest on-target activity in human cells (Supplementary figure 5). Because our major goal was to select Cas9 mutants without compromised on-target activity, we chose to characterize Sniper-1. Sniper-1, along with rationally designed Cas9 variants (eSpCas9-1.1 and Cas9-HF4; versions with highest specificities) and WT-Cas9, were tested in HEK293T cells with ten different targets. Because sgRNAs begin with guanine at the transcription start site to allow expression from U6 or T7 promoters, the targets could be divided into two groups (Supplementary figure 8): those with a mismatched guanine at the 5’ end (gX19 and gX20) and those with a matched guanine at this position (GX19). As reported by other groups^4–6^, with the gX19 and gX20 sgRNAs, eSpCas9-1.1 and Cas9-HF4 were poorly active, showing a median of 10% and 3%, respectively, of WT-Cas9 activities, whereas Sniper-1 maintained high activity levels comparable to WT-Cas9 (Figure 1b). With the GX19 sgRNAs, eSpCas9-1.1 and Sniper-1 showed high activities, whereas Cas9-HF4 was not efficient, with an average relative activity of 55%.

We next compared the specificities of Sniper-1 with those of eSpCas9-1.1 and Cas9-HF4 by calculating the ratios of on-target activity to off-target activity. In this analysis, we excluded Cas9 variant-sgRNA combinations with less than 70% on-target activity relative to WT-Cas9 (Figure 1c, Supplementary figure 9). For the gX19 group, only one target (*DMD*) passed this criterion with eSpCas9-1.1, and none with Cas9-HF4. In this case, Sniper-1 showed 5.7-fold higher specificity than eSpCas9-1.1. When Sniper-1 was combined with GX19 or gX20 sgRNA and the specificities compared, the optimum performance was observed with gX20 sgRNA, which is in line with the fact that the Sniper-screen was performed with gX20 sgRNAs. For the GX19 group, all five targets passed the 70% criterion for Sniper-1 and eSpCas9-1.1, whereas only the *FANCF01* target passed for Cas9-HF4. When the specificity ratios were compared for the *FANCF01* target for the Cas9 variants in their optimum combinations, Sniper-1 showed slightly better specificities than eSpCas9-1.1 and Cas9-HF4. When Sniper-1 and eSpCas9-1.1 were tested in their optimum combinations, specificities at least 10-fold higher than that of WT were observed for both mutants except at the *EMX1* and *FANCF02* targets, at which Sniper-1 showed 3.6-fold and 1.2-fold higher specificities than eSpCas9-1.1. We tested Sniper-1 and the other Cas9 variants in Hela cells (Supplementary figure 10). Similar results were obtained, confirming that the high specificity of Sniper-1 was not cell line specific.

An alternative way of measuring the specificity is to test a series of guide RNAs containing mismatches relative to the on-target sequence (Figure 2a, Supplementary figure 11). Among three different targets tested, direct comparisons between all of the Cas9 variants and WT-Cas9 were possible only at the *FANCF01* target site owing to the low on-target activities of eSpCas9-1.1 and Cas9-HF4 (Supplementary figure 11). Sniper-1 exhibited the highest specificities, discriminating single-nucleotide mismatches in the sgRNAs at every nucleotide position except position 19 when compared to WT-Cas9, eSpCas9-1.1 and Cas9-HF4. Almost no off-target activity was observed with Sniper-1 combined with sgRNAs with double mismatches.

**Figure 2.**
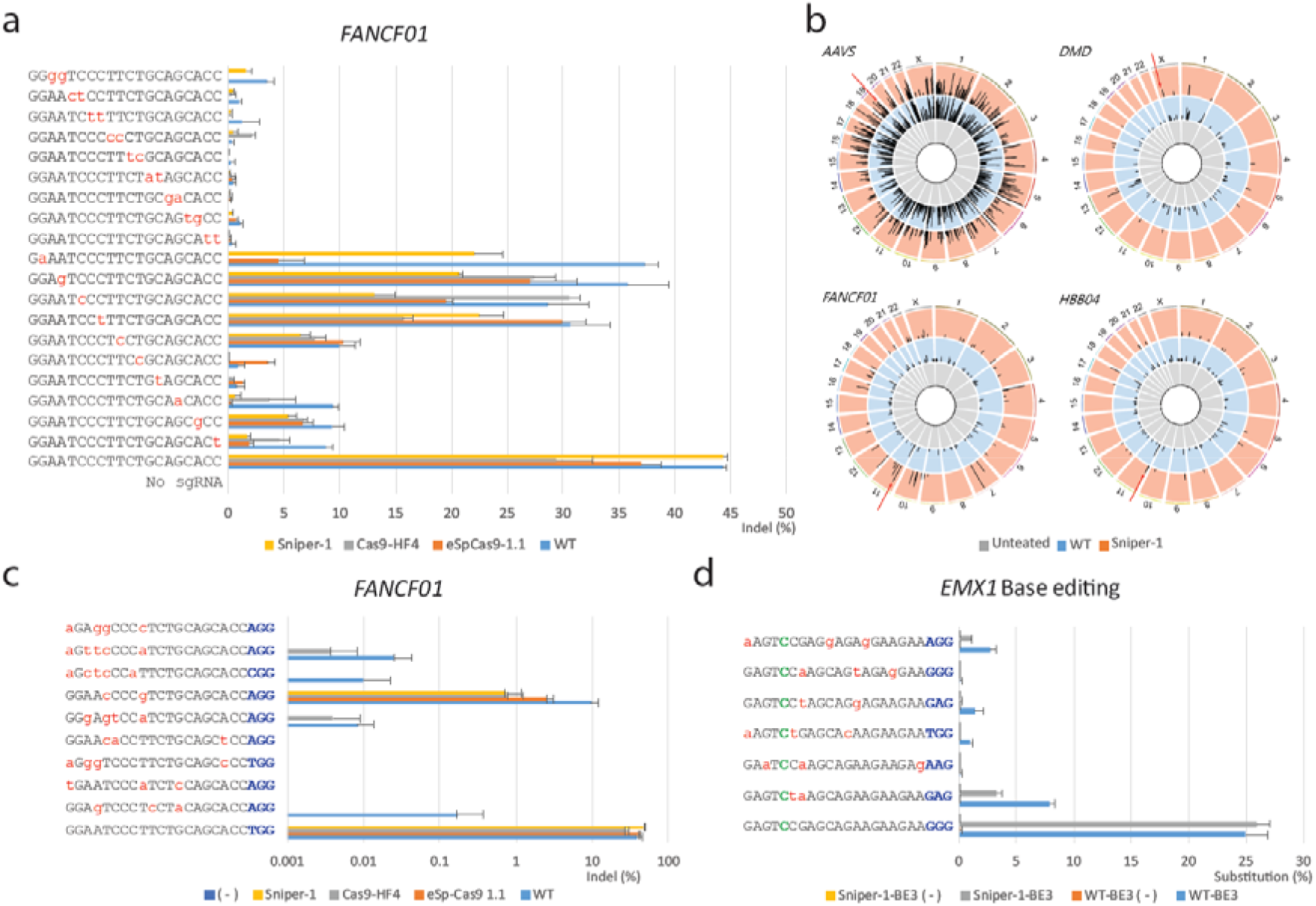
(**a**) Tolerance of Cas9 variants for sgRNAs with mismatches relative to the *FANCF01* target. Frequencies of small insertions or deletions (indels) were measured using targeted deep sequencing. Error bars indicate s.e.m. (*n*=3) (**b**) Genome-wide Circos plots representing DNA cleavage scores for *AAVS, DMD, FANCF01* and *HBB04* obtained with genomic DNA digested with untreated (grey), WT-Cas9 (blue) or Sniper-1 (orange). Arrows indicate on-target sites. (**c**) WT-Cas9, eSpCas9-1.1, Cas9-HF and Sniper-1 off-target sites for *FANCF01* validated in HEK293T cells by targeted deep sequencing. Error bars indicate s.e.m. (*n*=3) The PAM is shown in blue. (**d**) Base editor (BE3) on-target and off-target activities measured in HEK293t cells. (-) indicates the absence of sgRNA. Substitutions were measured using targeted deep sequencing. Substitution of C5 (represented by green type) to T was measured. The PAM is shown in blue. Error bars indicate s.e.m. (*n*=3)

Next, we proceeded to test the genome-wide specificity of Sniper-1 by performing multiplex Digenome-sequencing^12, 13^ ‘ with four different targets. The data showed significantly fewer (1.8~4.3-fold fewer) off-target candidates for Sniper-1 compared to WT-Cas9 (Figure 2b, Supplementary figure 12 and Supplementary figure 13). Ten candidates with the highest DNA cleavage scores were selected and validated by targeted deep sequencing (Figure 2c, Supplementary figure 14). Sniper-1 exhibited no measurable off-target mutations even with deep sequencing at all potential *FANCF01* off-target sites identified by Digenome-seq, except for one at which higher specificities were observed compared to WT (16.6-fold), eSpCas9-1.1 (3.92-fold) and Cas9-HF4 (1.84-fold). For the rest of the targets, a comparison was made between Sniper-1 and WT due to the low on-target activities of the other variants. Again, Sniper-1 showed much lower genome-wide off-target activity compared to WT. In particular, no off-target indels were measurably induced by Sniper-1 at Digenome-positive sites with two or more mismatches (Supplementary figure 4).

We also investigated whether the mutations in Sniper-1 can improve the specificity of base editors (BEs). To this end, Sniper-1 mutations were introduced into base editor 3 (BE3)^14^ to create Sniper-1 BE3, which was tested in HEK293T cells to determine its base-editing efficiency (Figure 2d). Sniper-1 BE3 was as efficient as WT-BE3 at the *EMX1* on-target site. At several pre-validated off-target sites that had been identified by Digenome-seq^15^, Sniper-1 BE3 showed much reduced off-target base editing effects (2.4~16.2 fold less) compared to WT-BE3. This result suggests that the nickase version of Sniper-1 also exhibits higher specificity than the WT Cas9 nickase, without sacrificing its on-target activity, unlike the nickase form derived from Cas9-HF (HF-BE3), which shows ~70% on-target activity compared to WT-BE3^16^.

In summary, we developed the Sniper-screen in E. coli to create a SpCas9 variant with increased specificity and full on-target activity. It is anticipated that further screening of Sniper-Cas9 or other Cas9 variants by the Sniper-screen would generate highly efficient and specific derivatives that could be used as therapeutic agents.

## Acknowledgments

This research was supported by grants from the Institute for Basic Science (IBS-R021-D1) to J.-S.K., Ministry of Science and ICT of Korea (2017M3A9B4061406) and Ministry of Agriculture, Food and Rural Affairs (MAFRA, grant number; 116088-3) to JKL. The plasmid encoding the pCMV-BE3 was a gift from David Liu (Addgene plasmid # 73021), pGRG36 was a gift from Nancy Craig (Addgene plasmid # 16666) and p11-LacY-wtx1 was a gift from Huimin Zhao.

## Author contributions

J.-S.K., S.K. and J.K.L supervised the research. J.K.L, E.J., J.L., M.J., E.S., Y.K. and K.L. performed the experiments. D.K. carried out bioinformatics analyses.

## Materials and Methods

### Plasmid construction

Each type of plasmid used in the Sniper-screen contains replication origins and resistance markers that are compatible with each other. (Supplementary Figure 2) The *ccdB* plasmid (p11-lacY-wtx1) was a kind gift from the Zhao lab^7^. It was double digested with SphI and XhoI enzymes (Enzynomics) which was ligated to oligos (Cosmogenetech) containing target sequences (Supplementary table 1) with T4 DNA ligase (Enzynomics). The sgRNA vector was constructed with a temperature-sensitive Psc101 replication origin^10^ (from pgrg36, a kind gift from Nancy Craig), tetR (from the tn10 locus of ElectroTen-Blue Electroporation Competent Cells, Agilent), a Kanamycin resistance marker, the pltetO1 promoter and the sgRNA sequence (synthesized at Bioneer). The components were PCR amplified and Gibson assembled. The guide RNA sequences to *EMX1* with various mismatches were cloned into the vector after BsaI digestion. The Cas9 library plasmid was derived from human codon optimized SpCas9 (Toolgen), dual CMV-pltetO1 (synthesized at Bioneer) and the p15a replication origin and chloramphenicol resistance marker (from the PBLC backbone, Bioneer). The components were Gibson assembled.

### *EMX1* genome insertion

Human *EMX1* containing various sgRNA target sequences (~500bp) was PCR amplified and integrated into the pgrg36^10^ vector between the NotI and XhoI sites. The cloned pgrg36-EMX1 vector was then transformed into the BW25141 strain to integrate the *EMX1* sequence into the tn7 site in the genomic DNA. EMX1-BW25141 was selected using the standard pgrg36 protocol^15^ (Supplementary Figure 1).

### Library construction

SpCas9 mutant libraries were constructed using three independent protocols. For the first library, the plasmid was transformed into XL1-red competent cells (Agilent), which were grown according to instructions in the vendor’s manual. For the second and third libraries, error prone PCR was performed on whole SpCAS9 sequences using Agilent and Clontech kits under low error-rate conditions. 3 × 10^6^ colonies were obtained for each library, resulting in a library complexity of 10^7^ overall.

### Positive and negative screening for evolving SpCas9 in E. coli

BW25141-EMX1 was co-transformed with the *ccdB* and sgRNA plasmids. The transformed BW25141-EMX1 cells were plated on ampicillin/kanamycin LB plates, which were then incubated overnight at 32°C. Transformants were cultured in liquid S.O.B. medium containing 0.1% glucose, ampicillin and kanamycin for electrocompetent cell production (Sniper-Screen). The SpCas9 library was transformed into the electrocompetent Sniper-Screen cells using a Gene Pulser (Gene Pulser II, Bio-Rad) following the manufacturer’s instructions. Transformed Sniper-screen cells were incubated in S.O.C. medium containing 10 ng/ml anhydrotetracycline (ATC) for 1hr at 37°C. The recovered Sniper-screen cells were plated on chloramphenicol/kanamycin LB plates (non-selective conditions) and chloramphenicol/kanamycin/arabinose LB plates (selective conditions) containing 10 to 100 ng/mL ATC followed by overnight culture at 32°C. Viable colonies were counted using OpenCFU^17^ and the survival frequency was calculated (survival frequency = the number of colonies on a selective plate/ the number of colonies on a non-selective plate). Colonies on the selective plate were pooled and incubated in chloramphenicol-containing LB medium overnight at 42°C to clear *ccdB* and sgRNA plasmids. Screened SpCas9 variant plasmids were purified using a midi prep kit (NucleoBond Xtra Midi EF, Macherey-Nagel) and continuously transformed into the Sniper-screen until the survival frequency reached a plateau. The concentration of ATC was gradually reduced to set harsher selection conditions using the process described above. Selected SpCas9 gene variants were shuffled to increase library diversity (DNA-Shuffling Kit, Jena Bioscience). Screening of the shuffled SpCas9 library was performed again and colonies on selective plates were individually cultured at 42° C to obtain evolved SpCas9 mutant plasmids. Each plasmid was Sanger sequenced using sequencing primers (Supplementary table) and three variants were chosen to be tested in a human cell line.

### Plasmids encoding Cas9 variants and sgRNAs

The WT-Cas9 encoding plasmid and U6 promoter-based sgRNA expression plasmid have been described previously^18^. Plasmids encoding human codon optimized eSpCas9 and Cas9-HF proteins were obtained from Addgene^2, 3^ and their Cas9 sequences were cloned into our WT-Cas9 backbone.

### Cell culture and transfection conditions

HEK293T cells (ATCC, CRL-11268) were maintained in DMEM medium supplemented with 10% FBS and 1% antibiotics. For WT-Cas9, eSpCas9-1.1, Cas9-HF4 and Sniper-1 mediated genome editing, HEK293T cells were seeded into 48-well plates at 70–80% confluency before transfection and transfected with Cas9 variant expression plasmids (250 ng) and sgRNA plasmids (250 ng) using lipofectamine 2000 (Invitrogen). For Base editing, HEK293T cells (1.5 × 10^5^) were seeded on 24-well plates and transfected at ~80% confluency with Base Editor plasmid (Addgene plasmid #73021) (1.5 μg) or Sniper-1-BE3 expression plasmid and sgRNA plasmid (500 ng) using Lipofectamine 2000 (Invitrogen). Genomic DNA was isolated with the DNeasy Blood & Tissue Kit (Qiagen) 72 h posttransfection.

### *In vitro* cleavage of genomic DNA

Genomic DNA was purified from HEK293T cells with a DNeasy Blood & Tissue Kit (Qiagen). Genomic DNA (10 μg) was incubated with Cas9 or Sniper1 protein (100 nM) and 4 sgRNAs (75 nM each) in a reaction volume of 1 mL for 8h at 37°C in a buffer (100 mM NaCl, 50 mM Tris-HCl, 10 mM MgCl2, 100 μg/mL BSA, at pH 7.9). Digested genomic DNA was treated with RNase A (50 μg/mL) to degrade sgRNAs and purified again with a DNeasy Blood & Tissue Kit (Qiagen).

### Whole genome and digenome sequencing

Genomic DNA (1 μg) was fragmented to the 400-to 500-bp range using the Covaris system (Life Technologies) and blunt-ended using End Repair Mix (Thermo Fischer). Fragmented DNA was ligated with adapters to produce libraries, which were then subjected to whole genome sequencing (WGS) using a HiSeq X Ten Sequencer (Illumina) at Macrogen. WGS was performed at a sequencing depth of 30-40×. DNA cleavage sites were identified using Digenome 1.0 programs^13^.

### Targeted deep sequencing

Target sites and potential off-target sites were analyzed by targeted deep sequencing. Deep sequencing libraries were generated by PCR. TruSeq HT Dual Index primers were used to label each sample. Pooled libraries were subjected to paired-end sequencing using MiniSeq (Illumina).

